# The germline-restricted chromosome orchestrates germ cell development in passerine birds

**DOI:** 10.64898/2025.12.02.691941

**Authors:** Niki Vontzou, Yifan Pei, Israel Campo-Bes, Wolfgang Forstmeier, Moritz Hertel, Manuel Irimia, Bart Kempenaers, Sylvia Kuhn, Katrin Martin, Jakob C. Mueller, Kim Teltscher, Annelie Mollbrink, Xesús Abalo, Matthew T. Biegler, Simone Immler, Francisco J. Ruiz-Ruano, Alexander Suh

## Abstract

While the definition of germ cell fate has been extensively studied in model organisms, evolutionary innovations and mechanistic novelties may remain hidden in understudied systems. The phenomenon of programmed DNA elimination allows germ cells to acquire germline-restricted genes, offering a novel paradigm of germ cell specificity. In passerine birds, the germline-restricted chromosome (GRC) is eliminated from somatic cells in early embryogenesis, yet the role and consequences of its maintenance in the germ cells remain poorly understood. Here, using the zebra finch *Taeniopygia guttata* as a model, we combined RNA-seq and Spatial Transcriptomics to construct a high-resolution spatiotemporal expression map to understand the role of the GRC across germ cell development. We found a GRC-linked integrin-BMP signaling in maturing oocytes and *tfeb*_GRC_ upregulation at blastoderm embryos, suggesting the involvement of the GRC in oocyte maturation and germ cell determination. We also identified developmental specialization of GRC-linked gene expression relative to their paralogs on the autosomes and sex chromosomes, revealing a gene repertoire which promotes germline stemness and germline/soma distinction. Together, the passerine GRC constitutes a unique system that manifests germ cell complexity, whilst allowing pinpointing the effects on gene expression that may elucidate vertebrate germ cell fate.

## Introduction

The discovery of non-Mendelian inheritance across the Tree of Life has shed light on unexpected molecular and evolutionary mechanisms that open new perspectives on our understanding of complex traits (Ross 2024). Paradoxical segregation patterns leading to the elimination of genome fragments or entire chromosomes, which sometimes occur in pathological conditions, are also facilitated as part of the life cycle of some organisms through the phenomenon of programmed DNA elimination (PDE) (Wang and Davis 2014; Smith et al. 2021). In these organisms, chromosome fragments or entire chromosomes are excluded from the somatic cells during their germline/soma differentiation and the resulting germline-specific DNA is transmitted to the next generation (Dedukh and Krasikova 2022).

Among vertebrates, the ∼6,700 species of passerine birds represent the largest taxonomic group in which PDE has been described to date (Torgasheva et al. 2019; Borodin et al. 2022; Ruiz-Ruano et al. 2025). The first avian germline-restricted chromosome (GRC) was serendipitously found in the zebra finch (*Taeniopygia guttata*), while studying male and female meiosis (Pigozzi and Solari 1998; Borodin et al. 2022). The GRC is the largest chromosome in the zebra finch genome and is exclusively present in male and female germ cells, being absent from the cells of somatic tissues (Pigozzi and Solari 1998; Borodin et al. 2022). The zebra finch GRC is usually found in two copies in oocytes, forming a recombining bivalent, and in a single copy in spermatocytes (Pigozzi and Solari 2005; Torgasheva et al. 2019, 2021). In males, the presence of the GRC has been reported in primary spermatocytes, yet it is usually eliminated from male germ cells before the end of meiosis (Pigozzi and Solari 1998, 2005), although rare exceptions exist (Pei et al. 2022).

The GRC consists of paralogous sequences duplicated from the regular ‘A chromosomes’ (autosomes and sex chromosomes) whereby some of these paralogs are further amplified into many copies on the GRC (Itoh et al. 2009; Biederman et al. 2018; Kinsella et al. 2019; Asalone et al. 2021; Mueller et al. 2023; Schlebusch et al. 2023). Consequently, hundreds of paralogous loci have been annotated on GRCs (with some families massively amplified) (Ruiz-Ruano et al. 2025; Mueller et al. 2023; Schlebusch et al. 2023), spanning a spectrum from near-identical to strongly diverged relative to their A-chromosomal partners. This complex genomic architecture has posed significant challenges to generating GRC reference assemblies, initially restricting GRC research primarily to cytogenetic approaches (Torgasheva et al. 2019; Malinovskaya et al. 2020; Torgasheva et al. 2021; Sotelo-Muñoz et al. 2022). These studies have consistently shown that the GRC is absent from the soma of every individual songbird studied so far, implying a robust and stable mechanism of programmed DNA elimination during embryogenesis. Representing passerines which comprise approximately two-thirds of all bird species (Gill F et al. 2025), the zebra finch serves as a powerful model for studying primordial germ cell development and culture (Gessara et al. 2021; Jung et al. 2019; Biegler et al. 2025a). Using complementary long-read sequencing technologies, we recently assembled a zebra finch germline reference genome, providing the first detailed view of the structure and complexity of the GRC (Ruiz-Ruano et al. 2025). This new reference allows us to directly identify the effects of the GRC on gene expression during germ cell development, elucidating potential GRC-specific roles in the germ cell fate.

The study of early embryonic development in birds is challenging due to the practical difficulties in obtaining and studying pre-ovipositional intrauterine embryos, which are characterized by rapid meroblastic cleavage within the first twenty-four hours after fertilization (Sheng 2014; Rengaraj et al. 2020). As a result, avian germ cell specification remained poorly understood. In 2016, it was shown that the germline-specific RNA marker DAZL (Deleted in Azoospermia Like) is asymmetrically localized in the domestic chicken (*Gallus domesticus)* oocyte and zygote, suggesting that the germ cell specification relies on pre-determining maternal components–the so called ‘germ plasm’ (Lee et al. 2016; Kim and Han 2018). More recently, the germ plasm organizer, Bucky ball, was identified in chicken oocytes co-localizing with known germline markers (Klein et al. 2024). However, the mechanisms by which germline signaling becomes restricted to primordial germ cells (PGCs) and negatively regulated in somatic cells remain unclear.

The passerine GRC could provide a unique system of irreversible DNA-based germ cell determination, constantly restricting germline signaling to PGCs, thereby allowing the somatic lineage to undergo further differentiation. Notably, compared to chicken, zebra finch PGCs are more abundant in the early embryo and highly express germline RNA markers (Jung et al. 2019). The initial stage of PGC migration occurs during the primitive streak formation, where PGCs passively relocate to the anterior extra-embryonic region and are subsequently actively incorporated into the germinal crescent (Wakely et al. 1997; Kang et al. 2015). Zebra finch PGCs migrate to the germinal crescent at later stages compared to chicken PGCs (Jung et al. 2019). Such heterochrony in PGC differentiation onset was recently documented in embryonic gonads of zebra finches compared to chicken (Biegler et al. 2025b). These findings suggest key differences in germ cell specification and development in zebra finches relative to chicken, questioning the extrapolation of chicken germ cell fate decisions to passerines.

In this study, using RNA-seq and Spatial Transcriptomics, we generated a high-resolution gene expression map of GRC-linked genes and their A-chromosomal paralogs throughout zebra finch germline development to explore GRC functions and the effects of PDE in shaping germline complexity. In particular, we sought for maternal mRNAs in preovulatory oocytes and early cleavage embryos that could influence PGC specification. We identified key GRC-linked transcripts such as *itga11*_GRC_ and *bmp15*_GRC_ to be carried over from maternal oocytes to early cleavage embryos, representing potential candidate components of the germ plasm. We further pinpointed the expression of the A-chromosomal *rp1l1* and the *znf239* zinc-finger protein to be among the first genes activated following the initial wave of zygotic genome activation (ZGA), potentially mediating the transition to the second major wave of ZGA. During ovipositional embryonic development, we observed a specialized GRC-linked upregulation of the transcription factor *tfeb*_GRC_, suggesting a role in maintaining pluripotency of blastodermal zebra finch embryos. Next, we tracked the upregulation of GRC-linked expression in gonads of hatching and post-hatching chicks, as well as a subsequent more limited expression in adult gonads, showing highly dynamic GRC activity in primordial germ cells of the developing gonad. Furthermore, we highlight consistent germline-enriched expression of A-chromosomal *ago3* and *dnmt3a*, indicating potential germ cell-specific roles in (post-)transcriptional control guided by miRNA silencing and *de-novo* methylation, respectively.

Altogether, our findings reveal distinct GRC-driven transcriptional programs across key stages of germ cell development, depicting programmed DNA elimination and the GRC as a kaleidoscope for understanding germ cell complexity and modularity.

## Results

### Transcriptional roadmap of the zebra finch germline development

To profile the transcriptional landscape of a germline with germline-restricted genes and to identify the timing of GRC-linked expression with high resolution, we selected key stages of the zebra finch germline life cycle. In total, we produced RNA-seq libraries from 39 samples representing oocytes, embryos at different stages, hatchling and adult gonads (Figure 1A, Supplementary Table 1). We determined embryonic stages based on morphological criteria from Eyal-Giladi and Kochav for intrauterine to ovipositional stages and Hamilton and Hamburger for post-ovipositional stages (Eyal-Giladi and Kochav 1976; Hamburger and Hamilton 1951). Because we were interested in elucidating the potential role of the GRC in germline/soma distinction, we dissected mature oocytes from F1 follicles and intrauterine embryos (EGKII/IV) from the ovary and the female reproductive tract, respectively. To investigate transcriptional changes between a naive pluripotency and migratory state, we dissected ovipositional blastoderms at EGKVI/VIII and late gastrulas/early neurulas at HH5/6. Finally, to test for sex-biased expression in zebra finch gonads from newly hatched and adult birds, we dissected testes and ovaries on the day of hatching, four days after hatching and from 4-year-old adult birds in breeding condition (Figure 1A). We next processed all libraries with the nextflow nf-core pipeline, including read pre-processing and mapping against the reference zebra finch somatic genome concatenated with a GRC assembly from a different population (Rhie et al. 2021; Ruiz-Ruano et al. 2025; Forstmeier et al. 2007). We then performed a Principle Component Analysis (PCA) using all expressed genes and plotted the first two principal components and a clustered heatmap with pairwise Euclidean distances between the libraries (Supplementary Figure 1A-B). Both PCA and pairwise Euclidean distance clustering suggested seven main groups on the basis of their developmental stage and tissue of origin (Supplementary Figure 1A-B). Both the PCA and the correlation analysis did not suggest any separation between day 1 and day 4 gonads in either sex. Likewise, there was no clear separation between mature oocytes and intrauterine embryos (Supplementary Figure 1A-B).

**Figure 1:**
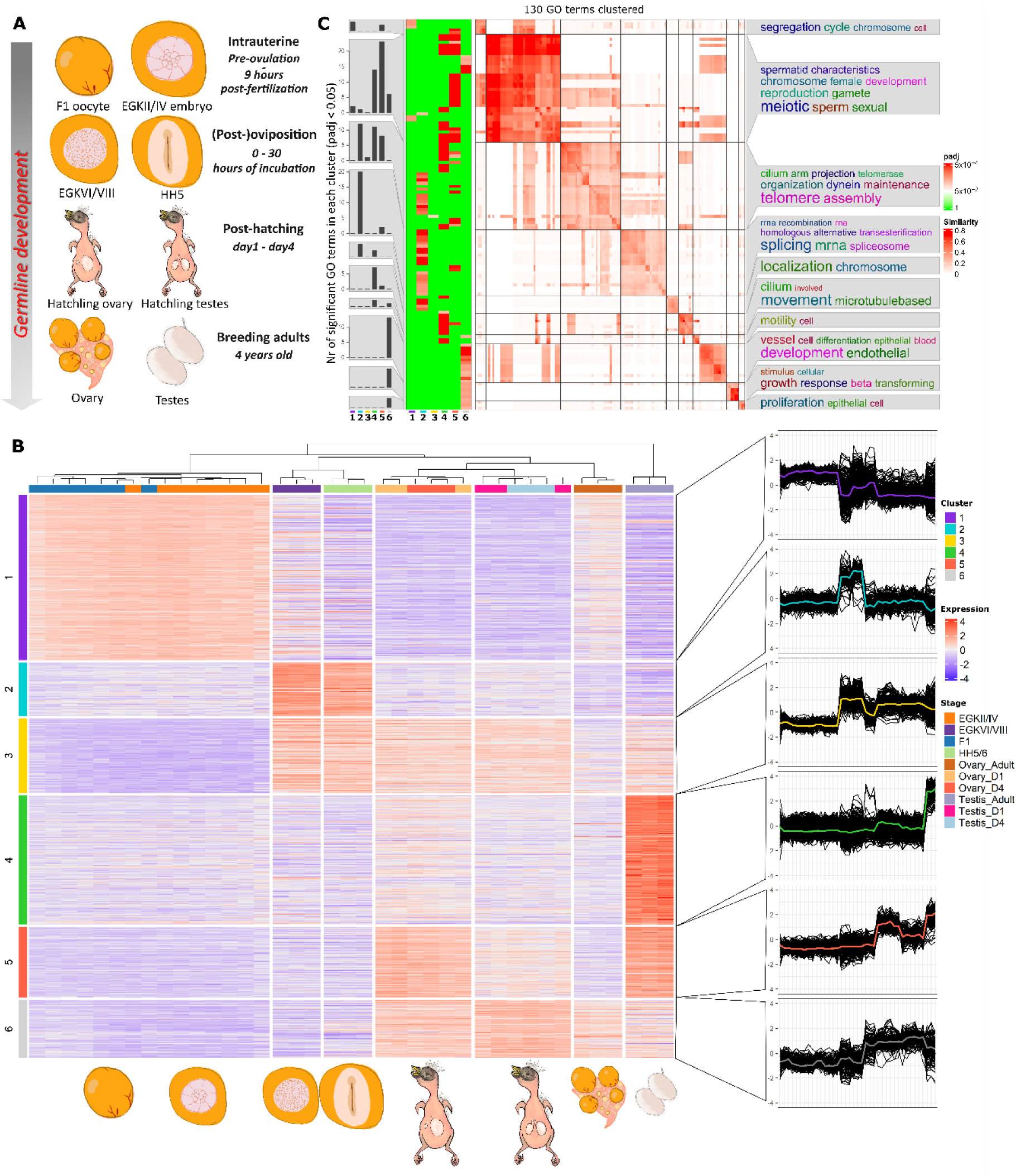
Developmental timing of zebra finch germline gene expression. A) Schematic illustration of the sampling stages for transcriptomic analysis, including preovulatory oocytes from F1 follicles, intrauterine EGKII/IV embryos within 9 hours after fertilization, EGKVI/VIII embryos from freshly laid eggs, HH5 embryos from 30 hour incubated eggs, gonads from day 1 and day 4 after hatching and 4-year-old male and female birds. B) Expression levels of the clustered top 10% of statistically significant genes detected by a likelihood ratio test (LRT) (FDR<0.001). The six gene clusters were generated by applying *k*-means clustering on rows of the expression matrix. The line charts (right) show the median expression for each gene cluster. The x-axis of each line chart is the same as the heatmap representing all samples per stage. C) Comparison of the enrichment results from over-representation analysis (ORA) performed on each gene cluster. Significant GO terms (p_adj_ <0.01) from the ORA of the six gene lists were clustered and their similarities were visualized (see Methods). The left heatmap shows whether GO terms were significant in the corresponding clusters. The middle heatmap represents similarities of clustered significant GO terms in any enrichment results. Font size of the word clouds (right) corresponds to the significance of the enrichment results.

Next, we performed a likelihood ratio test (LRT) to identify differentially expressed genes (DEGs) across all developmental stages, and we applied k-means clustering on the basis of the expression status of the top 10% significant genes (FDR < 0.001). The clustering analysis suggested six gene clusters with distinct expression patterns across development (Figure 1B). To identify enrichment changes across developmental stages, we applied an over-representation analysis (ORA) separately to the six gene clusters using zebra finch annotated genes as a background, and subsequently clustered their enrichment results (Figure 1C). We found gene clusters 1, 2 and 4 to be specifically upregulated in intrauterine and post-ovipositional embryos and testes, respectively. Cluster 3 comprised genes upregulated across both (post-)ovipositional embryonic and hatchling gonad development. Cluster 5 included genes upregulated in both hatchling ovaries and adult testes. Finally, cluster 6 entailed genes upregulated in all hatchling gonads, with some showing mild upregulation in adult ovaries.

Because paralogs of GRC-linked protein-coding gene symbols have a range of GRC-to-A-chromosomal sequence similarities, read mapping can lead to both ambiguous and unambiguous GRC-vs A-chromosomal-linked gene expression. To identify GRC-linked paralogs particularly prone to mis-mapping, we mapped brain RNA-seq data as a proxy for somatic gene expression to the same reference used in our study (Supplementary Table 2). We selected brain tissue because of its overall high transcriptional activity and overlapping expression profiles with the germline (Kulkarni et al. 2020). GRC-linked genes with all exons covered by the brain RNA-seq data were considered ambiguous in our subsequent analyses and therefore excluded (Supplementary Table 2). As a result, we focused solely on paralogous genes that were not part of the mis-mapping test and explored GRC-linked expression as part of the broader analysis of expression patterns and functional enrichment across the developmental stages.

Interestingly, we identified GRC-linked genes in all clusters except for cluster 4, which was testis-specific and enriched for biological processes such as cilium and microtubule-based movement (Figure 1C, Supplementary Table 3). Cluster 1 contained three GRC-linked genes (*itga11*_GRC_, *bicc1*_GRC_, *bmp15*_GRC_) and over-representation analysis revealed significant enrichment for chromosome segregation and cell cycle–related functions, consistent with intrauterine-specific upregulation in pre-ovulatory oocytes and cleavage stages (Figure 1C). Cluster 2 included a single GRC-linked gene (*tfeb*_GRC_) and was enriched for mRNA splicing and telomere assembly, suggesting high transcriptional activity following the maternal-to-zygotic transition after oviposition. Cluster 5 contained two GRC-linked genes (*puf60*_GRC_, *rxrb*_GRC_) and showed enrichment for meiotic cell cycle progression, indicating that cluster 5 is meiosis-specific. Cluster 6, which included five GRC-linked genes (*rnf17*_GRC_, *lin54*_GRC_, *bckdk*_GRC_, *elavl4*_GRC_, *ccdc25*_GRC_), was significantly enriched for processes such as cell differentiation, response to TGF-beta stimulus and endothelial development, suggesting a potential enrichment in germ cell differentiation. Cluster 3 included 17 GRC-linked genes (Supplementary Table 3), which showed no significant enrichment for any biological process, when using the zebra finch genome as background. To address potential limitations due to incomplete annotation, we repeated the over-representation analysis using the better-annotated chicken genome as background, which revealed a single significant enrichment for negative regulation of neuron projection development—a term uniquely enriched in cluster 3 and not observed in any other cluster (Figure 1C). Given the suggested pleiotropy showcased by the commonality in gene networks underlying germline and nervous tissues, this cluster of genes may facilitate the suppression of somatic genes in a germline-specific context and the promotion or/and maintenance of the germ cell fate (Kinsella et al. 2019; Kulkarni et al. 2020; Schlebusch et al. 2023). Of note, primordial germ cell regulators *dnd1* and *lin28* were found in cluster 3 and have been shown to suppress somatic differentiation of PGCs and promote PGC maintenance, respectively (Gross-Thebing et al. 2017; Suzuki et al. 2023).

### Transcriptome profiling of oocyte-to-embryo transition reveals key maternal components and early zygotic transcription preceding major genome activation

While F1 oocytes and intrauterine EGKII/IV embryos showed some overlap in the PCA (Supplementary Figure 1A, 2A-B), the EGKVI/VIII embryos formed a clearly distinct cluster, potentially indicating that zygotic genome activation (ZGA) has started but is incomplete in EGKII/IV embryos (Supplementary Figure 2C-D). This was also supported by the prominent elevation of intron-exon ratio for EGKVI/VIII relative to F1 oocytes, indicating a higher increase in nascent RNA synthesis during EGKVI/VIII compared to EGKII/IV (Supplementary Figure 2C). The gradual ZGA onset in the zebra finch embryos is in accordance with the previously reported chicken ZGA taking place in two waves, a minor one at early cleavage stages (EGKII/IV) and a major one at late cleavage stages (EGKIV/VI) (Hwang et al. 2018; Rengaraj et al. 2020).

To further investigate the onset of ZGA in zebra finch, we conducted pairwise comparisons of gene expression levels among F1 oocytes, EGKII/IV and EGKVI/VIII embryos (Figure 2, Supplementary Table 4). The comparison of F1 oocytes and EGKII/IV embryos revealed downregulation of 462 genes in EGKII/IV embryos, while only 17 genes were upregulated (absolute *Log_2_FoldChange* >1, *p_adj_*< 0.01), suggesting initiated maternal RNA clearance and early gene activation in EGKII/IV embryos (Figure 2A, Supplementary Table 4-5). Among those 17 upregulated genes in EGKII/IV embryos, we identified *rp1l1* as the top significant gene, which encodes a member of the doublecortin family, regulating microtubule polymerization (Figure 2A) (Reiner et al. 2006; Liu et al. 2024). Following *rp1l1*, we detected upregulation of *znf239* (*loc116807117*), which encodes a KRAB box transcription factor that belongs to the krueppel C2H2-type zinc-finger protein family and has been annotated in 47 copies in the zebra finch somatic reference genome (Ecco et al. 2017; Rhie et al. 2021). Therefore, the upregulation of these two genes in EGKII/IV embryos points towards microtubule stabilization and transcriptional regulation by *rp1l1* and *znf239*, respectively, shortly before the second wave of zebra finch ZGA.

**Figure 2.**
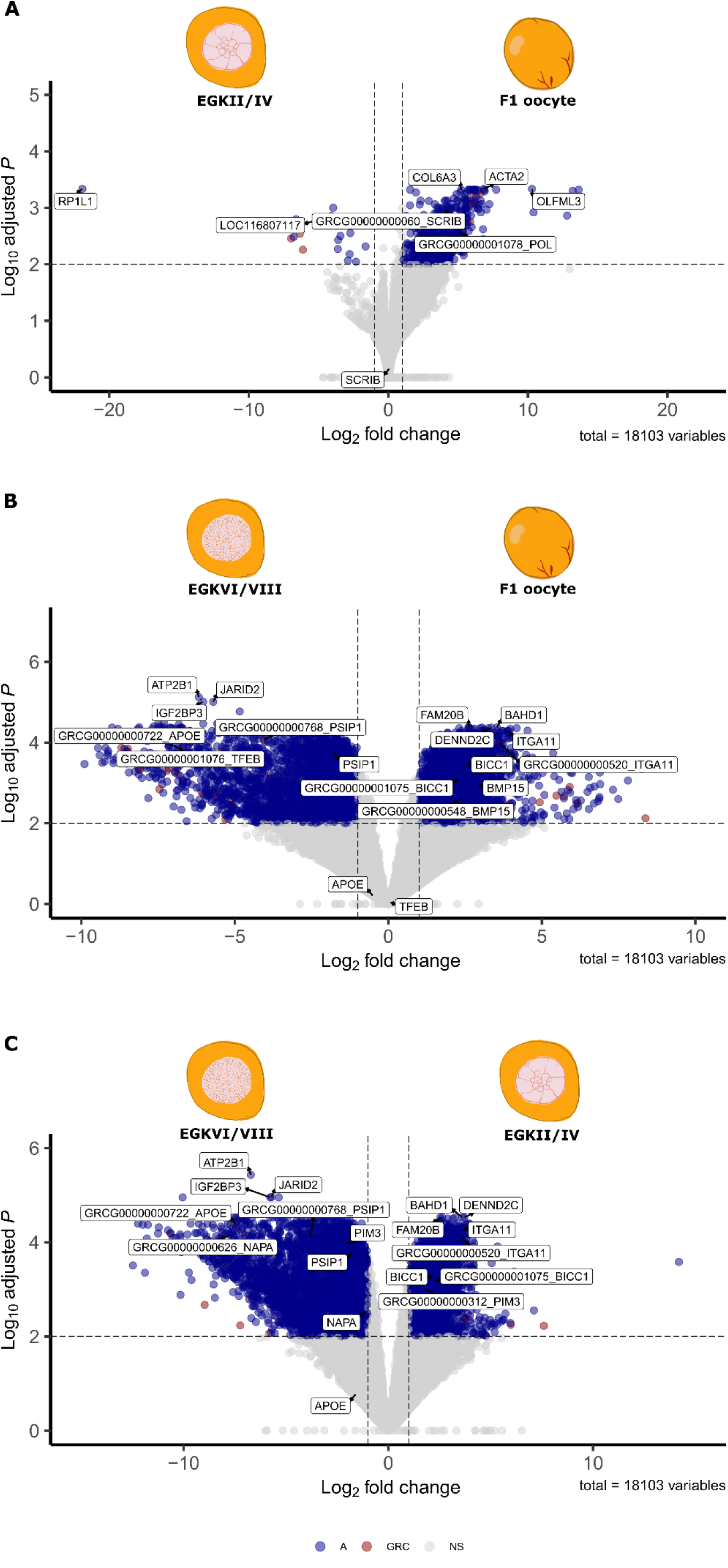
Maternal and zygotic genes in F1 oocytes, EGKII/IV and EGKVI/VIII embryos. A-C) Differential gene expression of the three pairwise comparisons between F1 oocytes, EGKII/IV and EGKVI/VIII embryos using DESeq2 (n = 7, n = 8, n = 3 from distinct F1, EGKII/IV and EGKVI/VIII samples; p_adj_ <0.01 and log_2_foldchange (LFC) < −1 or LFC > 1). Top A-chromosomal and GRC-linked significant genes per stage are shown along with their respective paralogs if the latter exist.

Among the DEGs between F1 oocytes and EGKII/IV embryos, we identified 11 GRC-linked genes upregulated in F1 oocytes and six in EGKII/IV embryos (Supplementary Table 4). Of the GRC-linked genes, *scrib*_GRC_ and *pol*_GRC_ were the only functionally annotated gene symbols and were downregulated in EGKII/IV embryos (Figure 2A). *Scrib* encodes a scaffold protein involved in cell polarity (Bera and Loeffler 2025; Stephens et al. 2018) and given that its GRC paralog *scrib*_GRC_ showed significant downregulation in EGKII/IV embryos, it is likely that *scrib*_GRC_ has a potential role in the asymmetric distribution of maternal components in the zebra finch cleavage embryo. It is noteworthy that in contrast to *scrib*GRC, the A-chromosomal paralog *scrib*A did not display a statistically significant change in mRNA level (Figure 2A). Finally, *pol*_GRC_ encodes for a polymerase that might belong either to endogenous retroviruses (ERVs) or genes derived from ERVs.

Overall, we observed the highest number of significantly DEGs when we compared freshly laid EGKVI/VIII embryos with intrauterine F1 oocytes and EGKII/IV embryos (Supplementary Table 4,6,7). Notably, we observed a substantial overlap of significantly DEGs in F1 oocytes and EGKII/IV embryos (Supplementary Table 8). The most significant DEG in both comparisons with EGKVI/VIII was *bahd1*, which is involved in heterochromatin formation and transcriptional repression (Figure 2B,C) (Bierne et al. 2009; Fan et al. 2021). Its persistent upregulation from F1 oocytes to intrauterine embryos indicates a role in maintaining transcriptional silencing during the minor ZGA, followed by its downregulation and an increased global transcription during the post-major ZGA at EGKVI/VIII embryos. Concomitantly, we observed a general increase in GRC-linked expression in EGKVI/VIII embryos relative to F1 oocytes and EGKII/IV embryos, with 53 and 58 GRC-linked significantly upregulated genes, respectively (Supplementary Table 4). *itga11*_GRC_ was the most significant GRC-linked gene shared in both F1 oocytes and intrauterine embryos when compared to EGKVI/VIII embryos (Figure 2B,C). Integrins are key players in linking the extracellular matrix to the actin cytoskeleton (Tanentzapf et al. 2007; Kanatsu-Shinohara et al. 2008). Intriguingly, integrin signaling has been found to control the germ cell niche in pluripotent cells in both *Drosophila melanogaster* fruit flies and the house mouse *Mus musculus*, suggesting a potentially shared function in germline stem cell niche regulation (Kanatsu-Shinohara et al. 2008; Makhlouf et al. 2024).

### Spatial transcriptomics of the finch blastoderm reveals the transcriptional landscape of germline and somatic lineages

To further investigate the expression of GRC-linked genes in the zebra finch blastoderm, we produced Spatial Transcriptomics data with 10x Visium for an EGKVI/VII embryo dissected from a freshly laid egg. We analysed a cross-sectional area on the yolk side of an individual blastoderm, capturing blastodermal cells of the germinal disc around the edge with yolk in the center (Figure 3A, Supplementary Figure 3). After clustering all spatial spots on the basis of the expression of common pluripotency markers *nanog* (*loc100230680*) and *pou5f1*(*loc100228280*) (Silva et al. 2009; Huang et al. 2009), we identified two superclusters representing the yolk- and blastodermal cell-enriched spatial spots (Supplementary Figure 3). Extraction and further clustering of the blastodermal supercluster was conducted to computationally derive two robust blastodermal subclusters (Figure 3B). We compared gene expression levels from the two subclusters to detect DEGs, identifying transcripts specific or enriched for somatic and germ cells, respectively (Figure 3C). The absence of genes with expression demarcated by the subcluster boundaries is most likely due to the oligo-cell resolution of the technology and the actual lack of morphological distinction between epiblast and hypoblast in the zebra finch blastoderm (Figure 3C) (Mak et al. 2015). The germline subcluster was enriched for *ddx4*, which is a well-known germline marker, as well as activin receptor type-2A *acvr2a*, which is expressed in chicken PGCs (Hickford et al. 2011; Whyte et al. 2015). Moreover, we found that the germline-enriched subcluster expresses more naïve pluripotency markers (*tfap2c*, *fbxo15, nrob1*) and *de-novo* methyltransferase *dnmt3a* compared to the soma-enriched subcluster (Pastor et al. 2018; Kojima et al. 2021; Liao et al. 2015; Nichols and Smith 2009; Dura et al. 2022; Ramabadran et al. 2023) (Figure 3C). By contrast, the general pluripotency markers, *pou5f1* and *nanog,* and *de-novo* methyltransferase *dnmt3b* were expressed in both soma and germline subclusters. The gene *tfeb*_GRC_ was enriched in the germline subcluster and was the most expressed GRC-linked gene with substantial GRC-specific single-nucleotide polymorphisms (SNPs) at its 3’-end. By contrast, *tfeb*_A_ did not display any expression. We therefore propose *tfeb*_GRC_ as a high-confidence GRC-linked marker specific to germ cells of the zebra finch blastoderm. *tfeb* is known to be involved in the regulation of pluripotency transcriptional network in undifferentiated mouse embryonic stem cells (Tan et al. 2021). Intriguingly, one of the few soma-enriched transcripts was *cerc2* (*loc100223021*), which is a chromatin remodeler involved in neurulation and somite development in chicken embryonic development (Chen et al. 2010) (Figure 3D). It is therefore plausible that distinct factors may regulate somatic differentiation versus germ cell pluripotency at this stage. The GRC may thus have a specific role in retaining a discrete pluripotency state in the early-specified primordial germ cells. Over-representation analysis showed a significant enrichment for post-translational modifications in the somatic subcluster, a process known to play key roles in embryonic stem cell pluripotency and differentiation (Figure 3E) (Wang et al. 2014). By contrast, viral processes were enriched in the germline subcluster, which may reflect pathways involved in the suppression of endogenous retroviruses (ERVs) (Prudhomme et al. 2005; Xiang et al. 2022) (Figure 3E). *Argonaute 3 (ago3)* was also significantly enriched in the germline subcluster, suggesting activation of an *ago3*-dependent miRNA-silencing pathway. Strikingly, *tnrc6a*, a core scaffolding protein of the miRNA-induced silencing complex (miRISC), was among the most upregulated genes in the somatic subcluster (Welte et al. 2023; Lazzaretti et al. 2009; Johnson et al. 2022). While *tnrc6a* is a well-known component of miRNA-mediated silencing, recent findings suggest it may also participate in additional mRNA silencing pathways through scaffolds with RNA-binding proteins (Welte et al. 2023). The distinct enrichment patterns of *ago3* and *tnrc6a* in germline and somatic subclusters, respectively, point to soma- and germline-specific engagement of distinct mRNA-silencing components, likely reflecting different recruiting mechanisms in post-transcriptional regulation. Together, these results reveal the gene expression dynamics underlying germline and somatic lineage specification, partially shaped by the presence of an irreversible germline-restricted DNA determinant.

**Figure 3.**
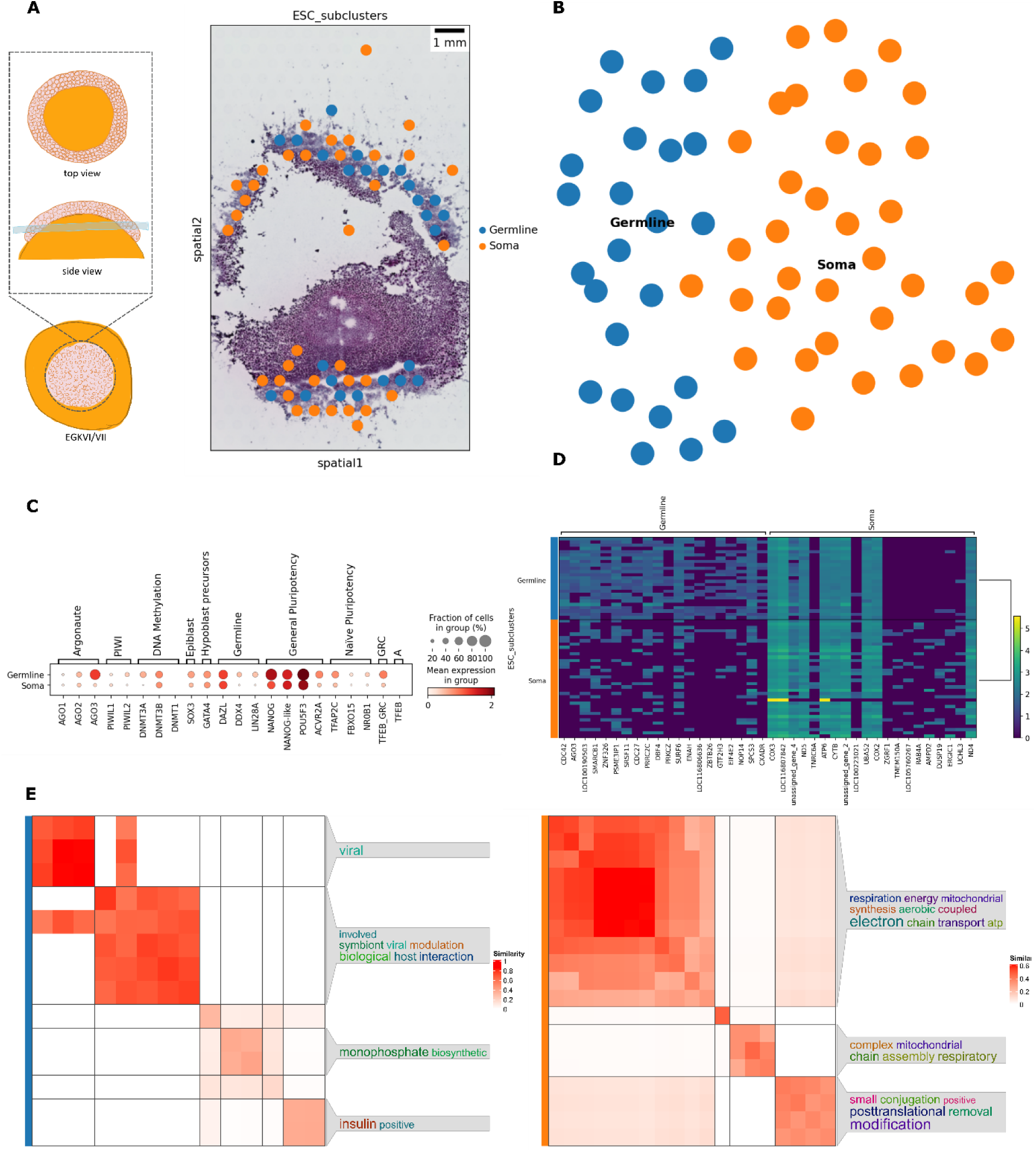
Spatial Transcriptomics of blastoderm EGKVI/VII zebra finch embryo. A) Top and side view of the embryonic cross-section capturing yolk in the middle and embryonic cells in the periphery. Arrowed boxes represent illustrations of the cross-section viewed from the horizontal plane (top) and the sagittal plane (side) (left). H&E (Hematoxylin and Eosin) stained histological section processed for Spatial Transcriptomics (right). B) Uniform Manifold Approximation and Projection (UMAP)-based dimensionality reduction plot of 10x Visium log-normalized data from a blastodermal horizontal section (blue: germline-enriched cluster, orange: soma-enriched cluster). C) Dot plot of mean expression values of marker genes for each cluster (color intensity denotes the mean expression of each gene in the given cluster; dot size denotes the percentage of spots in the given cluster expressing each gene). D) Heatmap with expression levels of top 20 marker genes across the two clusters (based on Wilcoxon rank test). E) Clustered significant GO terms enriched in each cluster (germline-enriched cluster on the left and soma-enriched cluster on the right) as predicted by an over-representation test of the top 100 DEGs of each cluster (FDR < 0.01).

### Expression profiles of GRC- and A-chromosomal paralogs suggest developmental specialization of GRC-linked genes

We investigated how the GRC-linked expression profile shifts during the PGC migration into the anterior extraembryonic region known as ‘germinal crescent’ (stage HH5-6). Strikingly, we found no differentially upregulated GRC-linked genes in the HH5 embryos (Figure 4A, Supplementary Table 9). By contrast, we identified several downregulated GRC-linked genes, with *tfeb*_GRC_ showing the most statistically significant downregulation, consistent with the EGKVI/VIII *tfeb*_GRC_ expression profile tracked by Spatial Transcriptomics (Figure 3C). Similarly, Spatial Transcriptomics in an HH5 embryo showed low GRC-linked gene expression, which may be partly caused by ambiguous read mapping from A-chromosomal paralogs (Supplementary Figure 4). Although we did not discern any GRC-linked genes with unambiguous read mapping in HH5 embryos, the spatial spots with the highest proportion in GRC-linked expression belonged to the hypoblast cluster, along with the most *dazl* and *ddx4* germ cell-specific expression (Supplementary Figure 4C). In the chicken embryo, PGCs ingress into the hypoblast and migrate to the germinal crescent through morphogenetic movements of the hypoblast (Kang et al. 2015; Ginsburg and Eyal-Giladi 1986). Here, the presence of germ cell markers in the hypoblast cluster most likely reflects the PGC-hypoblast adhesion during their migration to the anterior region (germinal crescent). The lack of GRC-linked expression in the HH5 embryos, as captured by both RNA-seq and Spatial Transcriptomics, could be in line with the transcriptional quiescence of PGCs after their specification to prevent their differentiation into somatic cell fates, previously reported in multiple vertebrates (Chihara and Nance 2012; Lebedeva et al. 2018; Robert et al. 2015).

**Figure 4.**
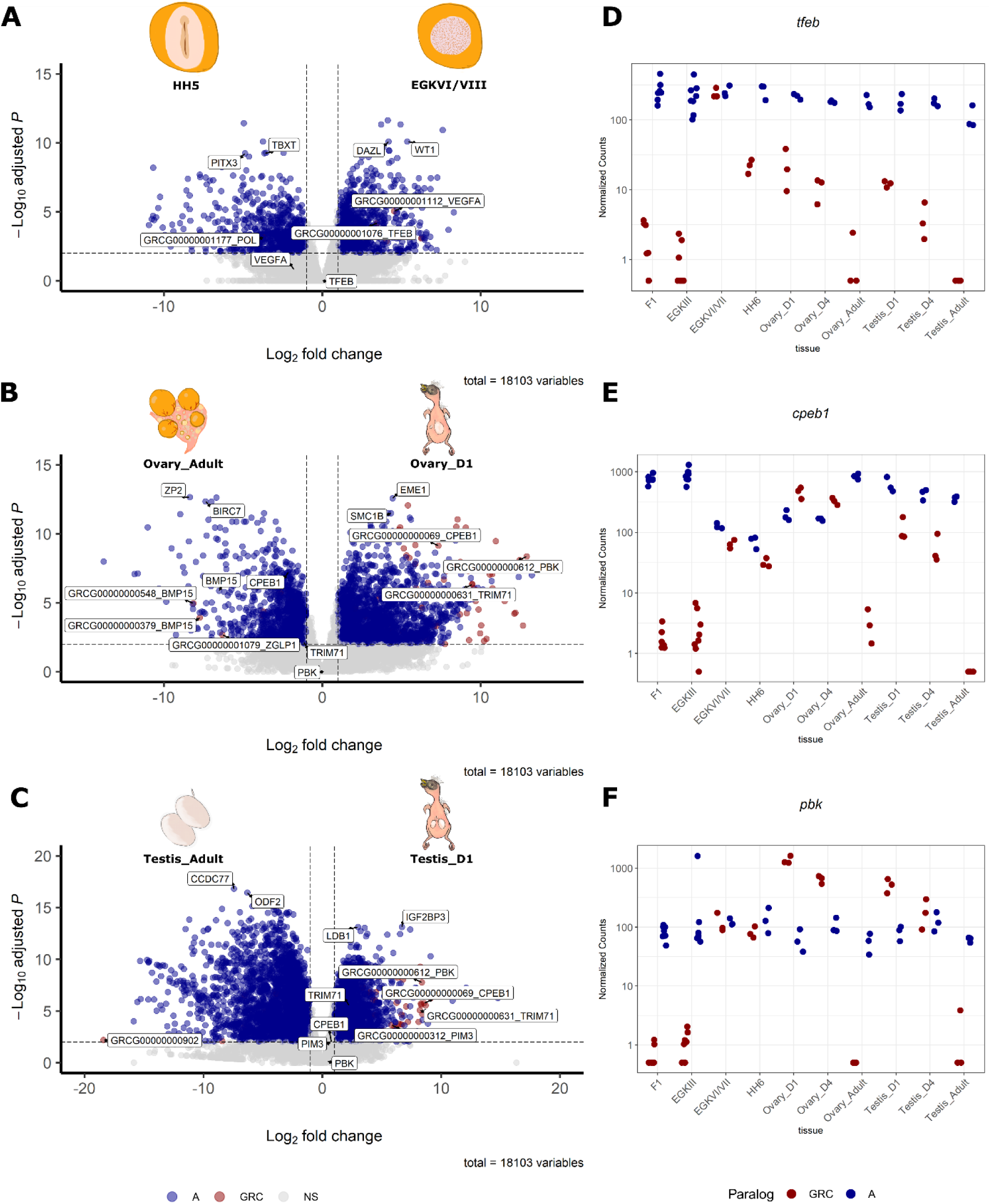
Specialized expression patterns of GRC-linked genes during embryo and gonad development. Differential gene expression of pairwise comparisons between EGKVI/VIII ovipositional vs HH5 incubated embryos (A), day 1 vs adult ovaries (B), day 1 vs adult testes (C) using DESeq2 (n = 3 replicates from distinct samples; p_adj_ <0.01 and log2foldchange (LFC) < −1 or LFC > 1). D-F) Normalized counts of selected genes across all replicates from each developmental stage (dark blue and dark red represents A-chromosomal and GRC-linked paralogs, respectively).

Next, to assess developmental gene expression differences in developing gonads, we conducted a comparative analysis of transcriptomic profiles between hatchling and adult gonads in both male and female individuals (Supplementary Table 10-11). It is important to note that the onset of female meiosis in chicken occurs asynchronously in late embryogenesis and gradually progresses upon hatching until meiotic arrest in prophase I (Smith et al. 2008). Interestingly, it has been previously shown that female proliferating germ cells were present in ovarian cysts on the day of chicken hatching (Yang et al. 2018). On the other hand, most male germ cells initiate mitotic arrest during embryogenesis, which persists upon chicken hatching, to only undergo meiosis during sexual maturation (Yang et al. 2018). Our differential expression analysis between hatchling and adult ovaries revealed a widespread downregulation of gene expression in adult compared to zebra finch day 1 ovaries (Figure 4B, Supplementary Table 10). Conversely, we observed the opposite pattern in zebra finch testes, with marked upregulation in adult testes (Figure 4C, Supplementary Table 11). These results conspicuously point to the aforementioned sexual dimorphism of gonad development on the day of avian hatching, driven by ongoing meiosis and active mitosis of female germ cells in contrast to the mitotically arrested male germ cells. Consistent with our clustering analysis (Supplementary Figure 1B, Figure 1B), few or no significant differences were observed between day 1 and day 4 zebra finch hatchlings in either sex (Supplementary Figure 5). This likely reflects pachytene progression in female gonads and sustained mitotic arrest in male gonads (Pigozzi and Solari 2005).

We observed an overall enrichment of GRC-linked DEGs being more expressed in day 1 gonads of both males and females (Figure 4B,C). *Pbk*_GRC_ and *cpeb1*_GRC_ were among the most upregulated GRC-linked genes in hatchling gonads of both sexes (Figure 4B,C). *Pbk* is a serine/threonine protein kinase and *cpeb1* is a cytoplasmic polyadenylation element binding protein; both are involved in cell cycle regulation (Dougherty et al. 2005; Rizkallah et al. 2015; Novoa et al. 2010). Notably, only few GRC-linked genes were significantly upregulated in the adult ovary. Among these, *bmp15*_GRC_ along with *zglp1*_GRC_ exhibited the highest GRC-linked expression (Figure 4B) (Paulini and Melo 2011; Rossetti et al. 2020). Of note, *bmp15* plays a crucial role in vertebrate oocyte maturation and folliculogenesis (Paulini and Melo 2011; Rossetti et al. 2020) and has 43 copy numbers on the zebra finch GRC (Ruiz-Ruano et al. 2025). Yet we only detected 8 copies with expression and among these *GRCG00000000548_BMP15*, *GRCG00000000379_BMP15* and *GRCG00000000732_BMP15* displayed the highest *bmp15*_GRC_ expression (Supplementary Figure 6A). *Zglp1* is the sole GRC-exclusive gene as its A-chromosomal paralog was lost in the common ancestor of ∼4000 bird species (Ruiz-Ruano et al. 2025) and interestingly is a key determinant of the oogenic cell fate in mice (Nagaoka et al. 2020). On the contrary, the differentially upregulated GRC-linked genes in the adult testis were either on the borderline of statistical significance or of unknown function (Figure 4C). These findings underscore the dynamic expression of GRC-linked genes during early gonad development and reveal a remarkably limited expression in adult gonads with the exception of *bmp15*_GRC_ and *zglp1*_GRC_ in the ovary.

We further explored the expression patterns of GRC-linked genes relative to their A-chromosomal paralogs across all selected developmental stages. *Tfeb*_GRC_ was strongly expressed in the zebra finch blastoderm of freshly laid eggs, as revealed by both RNA-seq and Spatial Transcriptomics. Its expression peaked at the embryonic stage EGKVI/VIII, while remaining low or undetectable at all other stages. By contrast, *tfeb*_A_ exhibited broad and consistent expression across all stages (Figure 4D). Similarly, *cpeb1*_GRC_ and *pbk*_GRC_ showed stage-specific upregulated expression, restricted to EGKVI/VIII and the hatchling gonads, while their A-chromosomal paralogs showed elevated expression trends across all stages (Figure 4E-F). These results suggest a more ‘specialized’, developmentally restricted expression pattern for some GRC-linked paralogs, while their A-chromosomal counterparts retain a developmentally broader expression (Figure 4D-F, Supplementary Figure 7). In contrast to this GRC-biased specialized expression across embryonic and gonad development, *bmp15*_GRC_ and *bmp15*_A_ paralogs displayed nearly identical expression profiles, since paralogs showed high expression in F1 oocytes, intrauterine EGKII/IV embryos, and adult ovaries (Supplementary Figure 7C). Notably, both *bmp15*_GRC_ and *bmp15*_A_ were upregulated in both F1 and EGKII/IV when compared to the later EGKVI/VIII stage, suggesting these are most likely maternal RNAs that are provisioned from oocytes to early cleavage embryos. These findings suggest that while several GRC-linked genes may have acquired specialized, stage-specific roles, others, such as *bmp15*_GRC_, retain developmental expression profiles similar to their A-chromosomal paralogs.

### Spatial Transcriptomics resolves folliculogenesis and spermatogenesis dynamics, revealing oocyte-specific GRC expression

Germ cell fate decisions are orchestrated in both time and space. To identify specific germ cell types of gametogenesis exhibiting GRC-linked gene expression, we performed Spatial Transcriptomics on adult gonads, both ovary and testis. Ovarian tissue was obtained from one sexually active female; following the dissection of the F1 oocyte for RNA-seq, the remaining ovary was subjected to histological sectioning and one cross-section was analyzed using 10x Visium Spatial Transcriptomics (Figure 5A). After performing clustering using the Leiden algorithm across a range of resolution parameters, we selected a resolution of 0.8, which provided optimal granularity for resolving both somatic and germ cell types. Our analysis yielded a total of 11 distinct cell clusters: three corresponding to germ cell clusters, six to ovarian somatic cell clusters, and two clusters belonged to the infundibulum of the oviduct (Figure 5B). We annotated the three germ cell clusters as early, late pre-hierarchical oocytes and ooplasm derived from the most mature F2 and F3 hierarchical oocytes of the sampled ovary (Apperson et al. 2017). In birds, folliculogenesis involves the sequential maturation of ovarian follicles from early, pre-hierarchical stages to fully developed hierarchical preovulatory follicles (Hlokoe et al. 2022; Kui et al. 2024). This process is tightly linked to meiotic progression: during folliculogenesis, oocytes resume meiosis from prophase I and progress to metaphase II, at which point they arrest again, shortly before ovulation (Rengaraj and Han 2022). We found both early and late pre-hierarchical oocytes expressing *zp2* glycoprotein subunit of the zona pellucida, while late pre-hierarchical oocytes further expressed A-chromosomal *cpeb1* and Aurora Kinase A (*aurka*). The co-expression of *cpeb1* and *aurka* in late pre-hierarchical oocytes (Supplementary Figure 8A) supports their coordinated role in meiotic progression, as Aurora Kinase A–mediated phosphorylation of CPEB1 is known to activate the translation of mRNAs critical for mouse oocyte maturation and meiotic progression (Kunitomi et al. 2024). Subsequently, the ooplasm from the maturing F2-3 oocytes was enriched for cyclin b2 (*ccnb2*), which has been previously found to be key for meiosis re-entry in mouse oocytes (Daldello et al. 2019). Additionally, *tacc3,* which has been proposed to play a role in stabilization of the mitotic spindle of mouse (Burgess et al. 2015; Ding et al. 2017), was upregulated in the F2-3 ooplasm cluster as well. Interestingly, we found *tacc3* to be further upregulated in both F1 and cleavage embryos when compared to the freshly laid blastoderms, indicating that the mRNA of this gene may be critical for the first divisions of the early embryo.

**Figure 5.**
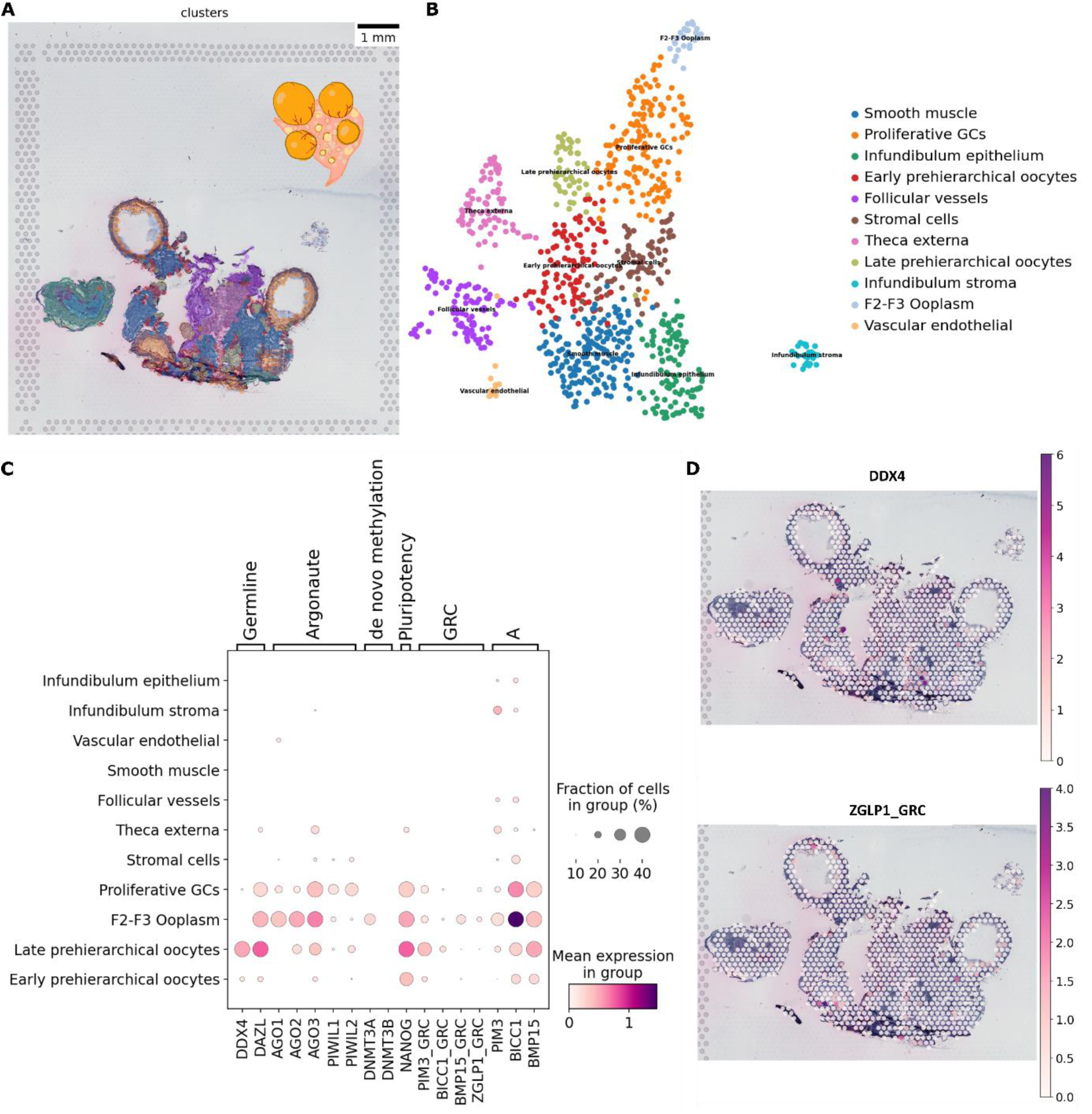
Spatial Transcriptomics of adult ovary in breeding conditions. A-B) UMAP plot of log-normalized clustered 10x Visium ovary data (n_pcs = 50, n_neighbors = 15, Leiden 0.8). C) Dot plot of mean expression of select marker genes across all clusters (Color intensity: mean expression level of the indicated gene over all spots in a given cluster; Size of dots: percentage of spots in specified cluster in which expression of the gene was detected). D) Scatter plot with overlay of expression values from germline marker gene *ddx4* and GRC-exclusive gene *zglp1GRC* on the hematoxylin and eosin stain (H&E) image.

The mRNAs of the germline markers *dazl* was predominantly detected in late pre-hierarchical and hierarchical oocytes and *ddx4* was expressed exclusively in late pre-hierarchical oocytes (Figure 5C). We also sought for the expression of argonaute genes as a proxy for the activity of RNA-guided RNA silencing pathways in female germ cells. We observed *ago2* and *ago3* to be highly expressed in the ooplasm of mature F2-3 oocytes, whereas *piwil1* and *piwil2* were primarily detected in proliferative granulosa cells (Figure 5C). These distinct expression patterns suggest active miRNA biogenesis in the ooplasm and piRNA-mediated silencing activity in the granulosa cells. To characterize the GRC-linked expression in the adult ovary, we focused on high-confidence GRC-linked paralogs with the highest transcript abundance across all spatial spots. *pim3*_GRC_ displayed expression in the late pre-hierarchical oocytes, whereas *pim3*_A_ was primarily expressed in the ooplasm of mature oocytes. Notably, *pim3*_GRC_-linked expression was consistently detected across all developmental stages examined, except in adult testes, suggesting a potential housekeeping role in the zebra finch germline (Ruiz-Ruano et al. 2025). By contrast, we detected *bmp15*_GRC_ and *zglp1*_GRC_ transcripts predominantly in the ooplasm of mature F2-3 oocytes (Figure 5C-D). The mature hierarchical oocytes are overall transcriptionally silent (Gaginskaya et al. 2009), suggesting that these genes are likely transcribed during earlier stages of oocyte maturation but are maintained in high levels in the cytoplasm until ovulation. We also detected the transcripts of *bmp15*_GRC_ and *zglp1*_GRC_ in EGKII/IV embryos during intrauterine development, indicating that the provisioning of these transcripts may be key for the early cell divisions in the early embryo (Supplementary Figure 7C-D). *Bmp15*_A_ was expressed in proliferative granulosa cells as well as in all three germ cell clusters and *zglp1*_GRC_ is the sole GRC-exclusive gene in zebra finch (Ruiz-Ruano et al. 2025). Together, these findings highlight an overall spatial and temporal specialization of specific GRC-linked genes in the female germline and suggest distinct roles during oocyte maturation and early embryogenesis.

The ability of Spatial Transcriptomics to simultaneously elucidate transcriptional changes across all cell lineages within the entire tissue (Moses and Pachter 2022) revealed the spatiotemporal dynamics of zebra finch spermatogenesis. We performed Spatial Transcriptomics on one cross-section each of the left and right testis from two individuals from two distinct captive zebra finch populations (Seewiesen and Krakow) (Forstmeier et al. 2007), in order to explore both intra- and inter-individual differences (Figure 6A). Upon integrating the left and right testis data for each individual, we observed convergent clusters, suggesting minimal intra-individual variation (Supplementary Figure 9). In contrast, when we integrated all datasets and repeated the analysis, we observed a post-meiotic population-specific distinction during spermiogenesis (Figure 6B-D).

**Figure 6.**
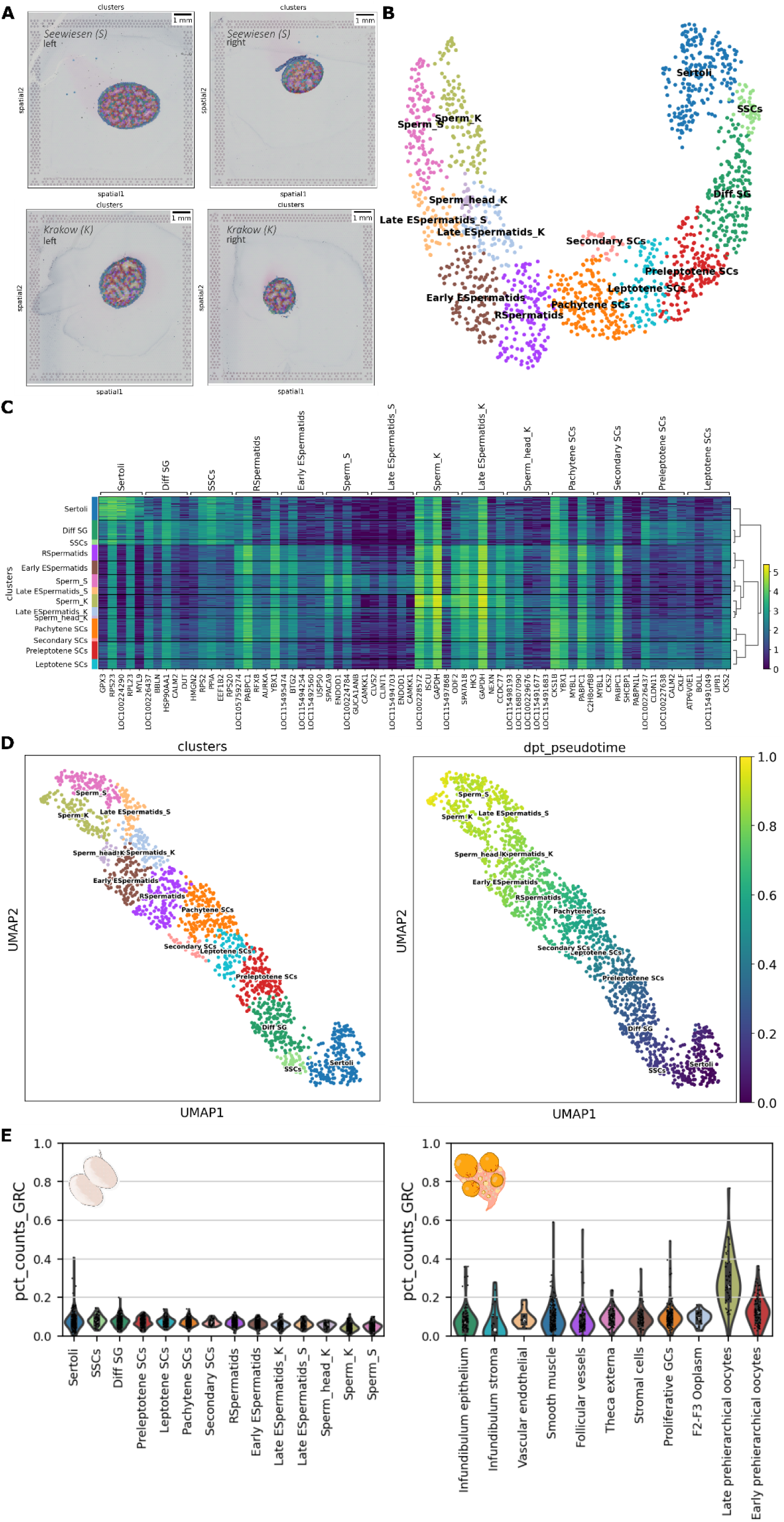
Spatial Transcriptomics in adult testes in breeding conditions. A-B) Log-normalized clustered data after integration of four testis 10x Visium data; left-right testis from a Seewiesen individual; left-right testis from a Krakow individual (Scanorma integration, n_pcs =30, Leiden 1.0). SSCs, Diff SG and SCs stand for spermatogonial stem cells, differentiated spermatogonia and spermatocytes, respectively. C) Heatmap of log-normalized expression values of the top 5 genes per cluster (Wilcoxon rank test). D) Diffusion pseudotime on a PAGA graph using Sertoli cluster as a root (UMAP was used to initialize the graph layout). E) Violin plots represent the percentage of GRC counts (pct_counts_GRC) across all clusters identified in integrated testis data and ovary data from Figure 5.

In contrast to the ovary, *Piwi* genes were more highly expressed than *Argonaute* genes in the testes, indicating activation of the piRNA pathway (Supplementary Figure 8B). *Piwil1* and *piwil2* were particularly strongly expressed in spermatogonia and primary spermatocytes, suggesting high piRNA activity in these cell types. Conversely, *dnmt3a* was enriched in both mature F2-F3 oocytes and multiple spermatogenic cell types, indicating a crucial role of DNMT3A-dependent DNA methylation in both oocyte development and spermatogenesis. GRC-linked expression was generally limited in the adult testes (Figure 6E), consistent with testis RNA-seq analysis and the absence of statistical power of GRC-linked expression in adults when compared to hatchling testes (Figure 4C, Supplementary Table 11), contrasting to our observation of the GRC-linked expression peaking in late pre-hierarchical oocytes (Figure 6E). The observed sex-biased GRC-linked expression pattern in adult gonads suggests absence of GRC testis-specific functions in adults and most importantly in spermatogonia, which is the most GRC-enriched cell type prior to GRC elimination during the first meiotic division in spermatocytes. The GRC-linked transcription in late-stage oocytes suggests a crucial role in oocyte maturation while the persistence of these oocyte-derived transcripts in early embryos implies a potential role during early cleavage stages.

## Discussion

The occurrence of a germline-restricted chromosome in the germline of a multicellular organism adds an additional layer to the germ cell mechanistic complexity, with potential effects on fundamental germ cell functions ensuring their dual capacity to self-renew and differentiate. In this study, using the zebra finch as model system of passerine GRCs, we presented a high-resolution roadmap of genome-wide gene expression across key stages of the germline life cycle, elucidating the roles of GRC-linked, A-chromosome-linked genes alongside their paralogs. Our wide and comprehensive sampling pinpointed key developmental time points with distinct GRC expression profiles reflecting GRC-specific roles in both self-renewal and differentiation of the germline. Beginning with the transcriptome of preovulatory oocytes, we reported the expression of genes involved in cell adhesion and polarity, namely *itga11*_GRC_ and *scrib*_GRC_. Given that avian germline/soma distinction follows the preformation model, characterized by asymmetric partitioning of maternal determinants during first embryonic divisions (Kim and Han 2018), we hypothesize that these GRC-linked transcripts may contribute to the specification of the germline niche during first cell divisions. Moreover, the expression of *bmp15*_GRC_ and *zglp1*_GRC_ in maturing oocytes points to their involvement in oocyte maturation before ovulation. *Bmp15*_GRC_ has repeatedly arisen across Passeriformes (Ruiz-Ruano et al. 2025), implying convergent evolution of germline-restricted *bmp15* paralogs and a selective advantage which may be associated with successful oocyte maturation and enhanced female fertility. *Zglp1_GRC_* is the sole GRC-exclusive gene detected in the common ancestor of ∼4,000 songbird species (Ruiz-Ruano et al. 2025) and its essential role in launching the oogenic program in zebra fish, mouse, and human oocytes (Wang et al. 2025; Nagaoka et al. 2020; Garcia-Alonso et al. 2022) implies a similar indispensable role in zebra finch oogenesis. We further demonstrated that transcriptional repression preceding the maternal-to-zygotic transition is likely mediated by *bahd1*, which is consistently expressed across all intrauterine developmental stages analyzed, suggesting that zygotic genome activation (ZGA) remains incomplete in EGKII/IV embryos. Notably, the persistence of *itga11*_GRC_, *bmp15*_GRC_, and *zglp1*_GRC_ mRNAs in EGKII/IV embryos indicates these are maternally inherited transcripts maintained throughout early mitotic divisions. Subsequently, during the minor wave of ZGA (Rengaraj et al. 2020), we observed the upregulation of microtubule stabilizer *rp1l1* and DNA-binding protein *znf239*, as well as GRC-linked genes of uncharacterized function. Overall, these findings reveal the earliest zygotically expressed genes in the zebra finch embryo, likely essential for the initial mitotic events, underscoring the key roles for both GRC-derived transcripts and A-chromosomal transcription factors that may orchestrate the transition to the major ZGA (Rengaraj et al. 2020). To the best of our knowledge, this is the first study providing insight into the intrauterine transcriptome of a passerine bird and the first characterization of the transcriptome of an animal before its developmental onset of PDE.

At the time of oviposition, we observed a repertoire of both GRC and A-chromosomal genes orchestrating soma and germ cell fate determination. Notably, we detected a markedly blastoderm-specific upregulation of *tfeb*_GRC_, while its A-chromosomal paralog *tfeb*_A_ showed consistent expression across all sampled stages as revealed by our RNA-seq data. Surprisingly, *tfeb*_A_ mRNA expression was not detected in the Spatial Transcriptomics data from the sampled region of the EGKVI/VII embryo, where epiblast-hypoblast segregation has not yet occurred, as indicated by the broad expression of hypoblast precursor *gata4* (Figure 3C). The absence of *tfeb*_A_ expression in the Spatial Transcriptomics data, in contrast to its presence in the RNA-seq of whole-mount embryos at the same stage, suggests that its expression may be restricted to certain regions, possibly in more differentiated hypoblast precursor cells, such as those in the central embryonic disc. This discrepancy could be due to the sampling limitations of the Spatial Transcriptomics approach, which may not have captured cells where *tfeb*_A_ is expressed. TFEB is a transcription factor that is well known for binding to lysosomal promoter target genes, yet cell-type dependent roles of TFEB outside the lysosomal pathway have been more recently reported. In human placental development, TFEB binds to promoters of syncytin genes and regulates their expression (Esbin et al. 2024). Notably, in mouse, *tfeb* has been previously shown to serve as a regulator of the pluripotency transcriptional network in embryonic stem cells, suggesting that *tfeb*_GRC_ may play a comparable role in maintaining pluripotency in the primordial germ cells of the zebra finch (Tan et al. 2021). Intriguingly, the zebra finch blastoderm at oviposition appears to be morphologically more similar to the mouse E4–E4.5 embryo and may exist in a more naïve pluripotent state compared to chicken (Mak et al. 2015). It is therefore intriguing to suggest that a more pronounced germline/soma determination, driven by the GRC, may be reflected by a prolonged naïve pluripotency state in the blastoderm zebra finch embryo. Our findings, highlighting the upregulation of both naïve pluripotency markers and *tfeb*_GRC_ expression within the germline-enriched cluster identified by Spatial Transcriptomics, further support this hypothesis.

Until now, gene expression analyses of the passerine GRC have been confined to adult gonads (Mueller et al. 2023; Kinsella et al. 2019; Schlebusch et al. 2023; Ruiz-Ruano et al. 2025). Our transcriptomic data reveal a significantly stronger enrichment of GRC-linked gene expression in both hatchling ovaries and testes relative to adults, underscoring a potential role for the GRC in gonad and PGC development. Consistent with this, striking differences in germ cell development between chicken and zebra finch embryonic gonads were recently reported (Biegler et al. 2025b). Although the expression of only few GRC-linked genes was examined, the presence of GRC-linked expression in PGCs of the zebra finch embryonic gonad suggested its involvement in these species-specific developmental programs (Biegler et al. 2025b). Importantly, our study demonstrated expression specialization (Mantica and Irimia 2025) of GRC-linked paralogs as indicated by lower expression breadth across developmental stages while A-chromosomal paralogs remain broadly expressed. For instance, we suggest that *tfeb*_GRC_ has acquired a specialized expression pattern at the time point of oviposition, while other GRC-linked genes, such as *cpeb1*_GRC_, *pbk*_GRC_ and *trim71*_GRC_, appear to be specialized across both (post-)ovipositional embryos and hatchling gonads (Figure 4E-F, Supplementary Figure 7). Consistent with this notion, we observed markedly reduced GRC-linked gene expression in adult gonads, with the notable exception of a subset of transcripts expressed during oocyte growth in the adult ovary, supporting an ovary-biased role of the GRC, especially in oocyte maturation. The strong signal of *bmp15*_GRC_ in maturing oocytes detected by both RNA-seq and Spatial Transcriptomics suggests that *bmp15*-dependent signaling is particularly critical during oocyte development (Supplementary Figure 6B). The expression of both *bmp15*_A_ and *bmp15*_GRC_ paralogs in zebra finch oocytes points to a dosage sensitivity, where an increased dose in *bmp15* may be required in regulating follicle growth, quality and subsequently female fertility (Supplementary Figure 6A-B) (Han et al. 2015; Persani et al. 2014). Notably, *bmp15GRC* has been previously highlighted to be among the most highly expressed GRC-linked genes in the blue tit ovary (Mueller et al. 2023), in agreement with the recently reported repeated duplications of *bmp15* from the A-chromosome onto the GRC across passerines (Ruiz-Ruano et al. 2025).

The existence of a pleiotropic relationship between germ cell and neural cell fates is underpinned by shared molecular mechanisms and gene expression networks between the two cell types (Kulkarni et al. 2020), and our study of GRC-linked genes further supports this observation. One of the most diverged genes in both zebra finch and nightingale germline-restricted chromosomes is *cpeb1*_GRC_ (Schlebusch et al. 2023) and CPEB1 is a key regulator of synaptic plasticity in vertebrate neurons (Chae et al. 2010). The reported specialized developmental expression patterns of GRC-linked genes, including *cpeb1*_GRC_, supports a potential role of the GRC in reinforcing developmental modularity of the germline-specific program. In this regard, we also highlight the identification of a distinct gene repertoire (Figure 1B-C, Supplementary Table 3), entailing both A-chromosomal and GRC-linked genes, consistently upregulated across primordial germ cell development and enriched for biological processes associated with the negative regulation of neuron projection development. Given the presence of pleiotropic genes arising from gene network co-option between germline and nervous system, the recruitment of a gene repertoire to promote germ cell fate while suppressing the somatic program may serve to enhance germ cell type specificity and tackle potential constraints imposed by pleiotropy (Kulkarni et al. 2020; Robert et al. 2015). Intriguingly, key germ cell factors such as *dnd1* and *lin28* as well as several GRC-linked genes are components of this repertoire, potentially reflecting an amalgam of A-chromosomal and GRC-linked genes responsible for maintenance of germ cell identity, while simultaneously suppressing the somatic program. The GRC could therefore allow the emergence of a “DNA-centered” germ cell-specific regulation, releasing potential constraints arising from a shared developmental program with the somatic cell lineage.

The role of the GRC-encoded protein toolkit in the maintenance of the germ cell fate and immortality is further underpinned by its developmental expression profile. The prominent expression of GRC-linked genes during embryonic and gonad development, along with their limited expression in adult gonads, supports a role in preserving the pluripotency of germ cells in the zebra finch. Consistently, we did not identify any GRC-linked genes within clusters enriched for meiotic processes, pointing towards a GRC-specific role in early germline specification and maintenance of germ cell identity, rather than gametogenesis. Intriguingly, genes with germline-specific expression that have been found to be expressed in human cancer, commonly referred to as germ cell cancer genes or testis/cancer genes, have been previously identified in passerine GRCs (Supplementary Table 12) (Vontzou et al. 2023; Kinsella et al. 2019; Ruiz-Ruano et al. 2025). For example, *tfeb* is an oncogene with a central role in the onset and development of various types of cancers, and our results of a specialized expression profile of *tfeb*<u>_GRC_</u> suggest a role in ovipositional primordial germ cell determination or/and maintenance (Supplementary Table 12). This observation resembles a more general phenomenon of cancer cells hijacking hallmarks of the germline to drive tumorigenesis, a process that is often described as the soma-to-germline transition (Bruggeman et al. 2023; Naik et al. 2024). PDE has been proposed to be capable of mitigating germline-soma conflicts by allowing the irreversible germline-specific expression of genes that may be detrimental for the soma (Smith et al. 2021; Vontzou et al. 2023), releasing the germline and soma from their conflicting interests imposed by antagonistic pleiotropy. Studying the complex gene network encoded by the passerine GRC may thus pinpoint genes that are exploited by cancer cells or/and underlie antagonistic pleiotropy.

Taken together, we present a high-resolution spatiotemporal gene expression map of programmed DNA elimination across the life cycle of the germline. Our findings highlight the potential roles of the GRC in germ cell determination, germ cell self-renewal and shaping specialized developmental expression profiles. Our approach thus opens up new directions for the study of GRCs and other forms of PDE, serving as a powerful model to illuminate the complexity of cell fate decisions, molecular interactions imposed by pleiotropy as well as the genetic and mechanistic basis shared between germline biology and cancer.

## Materials and methods

### Sample collection for RNA-seq and 10x Visium

For this study, we collected and dissected F1 oocytes and intrauterine EGKII-IV embryos from females with an ongoing clutch, as well as EGKVI/VIII and HH5 embryos from freshly laid and incubated eggs. All samples were obtained from breeding pairs of a domesticated zebra finch (*Taeniopygia guttata castanotis*) population (“Seewiesen”) maintained at the Max Planck Institute for Biological Intelligence since 2004 (Forstmeier et al. 2007). The only exception was the sampling of testes for Spatial Transcriptomics from a male individual from the captive “Krakow” population maintained at the Max Planck Institute for Biological Intelligence (Forstmeier et al. 2007). F1 oocytes and intrauterine embryos were collected simultaneously from the same breeding female. Dissections were performed during a consistent daily time window, and embryo staging was based on established morphological criteria (Hamburger and Hamilton 1951; Eyal-Giladi and Kochav 1976; Murray et al. 2013). Immediately after collection, F1 oocytes and intrauterine embryos were excised from the largest yolk-containing follicle and egg found in the reproductive tract, respectively, while EGKVI/VIII blastoderms and HH5 embryos were dissected from freshly laid and ∼30-hour incubated eggs in the nest, respectively. Samples were transferred to RNase-free phosphate-buffered saline (PBS), and yolk and vitelline membranes were removed following dissection of the embryo or oocyte.

Flash-frozen samples were collected for RNA-seq, and OCT-embedded snap-frozen samples were collected for 10x Genomics Visium Spatial Transcriptomics. Unfertilized or undeveloped embryos were identified based on their morphology and were subsequently excluded from sampling. Gonads were dissected from day 1 and day 4 hatchlings as well as from sexually mature four-year-old adults from the same Seewiesen population. Adult ovaries were sampled without the F1 oocyte and if the latter existed, it was sampled separately. Immediately after excision, gonads were either flash-frozen for RNA-seq or OCT-embedded and snap-frozen for Spatial Transcriptomics. All samples were stored at –80 °C prior to library preparation and sequencing. All animal procedures, including sacrifice and dissection, were conducted with the minimal required number of sacrificed birds under compliance with ethical best-practice and in accordance with permits (permit nos. 311.4-si and 311.5-gr, Landratsamt Starnberg, Germany).

### RNA isolation and RNA-seq

Total RNA from flash-frozen oocytes, embryos and hatchling gonads was isolated using the miRNeasy Tissue/Cells Advanced Micro Kit and total RNA from flash-frozen adult gonads was isolated using the miRNeasy Tissue/Cells Advanced Mini Kit (Qiagen) following the manufacturer’s indications. The quantity and quality of the extracted total RNA was determined using Qubit (Thermofisher) and Agilent 2100 Bioanalyzer (Agilent Technologies). Libraries were constructed with the SMARTer Total Stranded RNA-seq kit - Pico input mammalian including an rRNA depletion step and sequenced using the NovaSeq6000 platform, yielding an average read depth of 80 million reads per sample of 2 × 151 bp paired-end reads.

### Spatial Transcriptomics

Tissue optimization was first applied to establish the right permeabilization conditions for EGKVI/VII embryo, HH5 embryo, adult ovary and testis. In total, two slides for tissue optimization and two slides for library preparation were processed at SciLifeLab Stockholm. Permeabilization time for adult testis was 29 minutes whereas for embryos and adult ovary was 10 and 5 minutes, respectively, as a result two different library preparation slides were followed, one with embryos and adult ovary and a second one with adult testis. After cryosectioning the OCT-embedded samples, libraries were constructed using the Visium Spatial Gene Expression kit following manufacturer’s guideline (10x Genomics). Samples were sequenced on NextSeq2000 platform with a 28nt(Read1)-10nt(Index1)-10nt(Index2)-90nt(Read2) setup at SciLifeLab Stockholm, producing an average read depth of 100 million reads per capture area.

### Mis-mapping check

To identify GRC-linked genes prone to mis-mapping, we aligned RNA-seq data from zebra finch brain tissue to a custom reference genome that included the GRC assembly (zfGRC) (Rhie et al. 2021; Ruiz-Ruano et al. 2025). Specifically, raw reads from the brain RNA-seq library SRR16168753 (Lipshutz et al. 2022) were mapped to the same merged reference genome used in our downstream analyses, consisting of the bTaeGut1.4.pri primary genome assembly combined with the zfGRC assembly. Read alignment was performed using STAR (2.7.11a), followed by transcript assembly and quantification with StringTie (2.2.1) (Dobin et al. 2013; Pertea et al. 2015).

### RNA-seq analysis

All RNA-seq libraries were processed using the nf-core/rnaseq pipeline (v3.9) with default parameters and a custom configuration file (available in the supplementary materials) (Patel et al. 2025). Unique molecular identifiers (UMIs) were processed using the -- with_umi option, with extraction parameters --umitools_extract_method “regex” and --umitools_bc_pattern2 “^(?P<umi_1̾.{8})(?P<discard_1>.{6}).*”. Reads were mapped against the reference assembly described above using STAR aligner and Salmon quantification, as implemented in the pipeline (nf-core/rnaseq v3.9). Pairwise differential gene expression analysis was conducted using the R package DESeq2 (v1.46.0) with developmental stage as fixed effect (Love et al. 2014). P-values were calculated via Wald-test using the t-statistic by setting the parameter (UseT=True). Visualization of differential expression results with volcano plots was performed using R package EnhancedVolcano (v1.24.0) and specialization plots were made using ggplot2 (v3.5.2) (Hadley Wickham 2016).

### Clustering and gene set enrichment analysis

To analyze all developmental stages at once, a likelihood ratio test was applied using deseq2 (v 1.46.0) (Love et al. 2014). After rlog transformation and subsetting of the top 10 % of statistically significant genes (p_adj_< 0.001), k-means clustering was performed using pheatmap (Kolde 2019). Visualization of the clusters was done using the packages pheatmap (v1.0.12) and complexHeatmap (v2.22.0) and ggplot2 (v3.5.2) was used to plot linear charts of the expression per cluster (Kolde 2019; Gu et al. 2016; Hadley Wickham 2016). The R package BioMartGOGeneSets (v0.99.11) was used to obtain zebra finch and chicken gene sets supported in BioMart (Zuguang Gu 2023). Over-representation analysis (ORA) was applied using ClusterProfiler (v 4.14.6) for each cluster and the package simplifyEnrichment (v2.0.0) was used to compare and visualize the enrichment results across clusters (Yu et al. 2012; Gu and Hübschmann 2023).

### Spatial Transcriptomics analysis

Spatial Transcriptomics data for each tissue section were processed using SpaceRanger 2.0.1 with the reference genome assembly and annotation described above (bTaeGut1.4.pri + zfGRC) (Rhie et al. 2021; Ruiz-Ruano et al. 2025). All downstream analysis and visualization were conducted in Python using Scanpy (v1.9.1) and Squidpy (v1.2.2), within a reproducible conda environment (Wolf et al. 2018; Palla et al. 2022). Visium spot and gene filtering was applied based on the ngenes_by_counts x total_counts scatter plots per library. Normalization of each spot by total counts over all genes and log1p transformation was done using scanpy (counts_per_cell_after=1e4). UMAP-based dimensionality reduction (v0.5.3) and clustering with increasing levels of resolution was performed (Leiden 0.4-1.2) and the level of granularity was manually inspected. The final level of resolution was empirically selected based on optimal granularity and biological interpretability for each library. Cluster annotation was performed based on top ranked genes per cluster using a Wilcoxon rank test in combination with the expression of known markers from the literature. Integration of testis libraries was performed using scanorama, enabling cross-sample comparison (Hie et al. 2024). Trajectory inference was carried out using PAGA (Partition-based Graph Abstraction) as implemented in scanpy, and diffusion pseudotime was computed with a manually assigned progenitor cluster as the root (Wolf et al. 2019). Trajectories were visualized in UMAP space (v0.5.3).

## Data availability

Prior to publication of this manuscript, we will submit all sequencing libraries to the relevant public databases. Custom code will be made available on GitHub (https://github.com/nikivon).

## AI statement

This manuscript was written entirely by the authors’ natural intelligence.

## Acknowledgements

We thank the Suh Lab and the GRC Brainstorming group for all helpful discussions, Renuka Kudva (NGI Sweden) for all the insightful guidance on the RNA-seq experiments, the Gahr lab and especially Antje Bakker for all the tips on RNA extractions, and Andrea Munsterberg and Emily Smith for introducing us to avian embryo dissections. This work was funded by a Consolidator Grant of the European Research Council (101002158 GermlineChrom) to Alexander Suh, a SCI Postgraduate Research Fellowship from the University of East Anglia (UEA) to Niki Vontzou, a Project Grant of the Swedish Research Council Vetenskapsrådet (2020-04436) to Alexander Suh and Francisco J. Ruiz-Ruano, a Nilsson Ehle Endowment of the Royal Physiographic Society of Lund to Francisco J. Ruiz-Ruano, a Marie Skłodowska-Curie Individual Fellowship (875732 birdGRC) to Francisco J. Ruiz-Ruano, a Bioinformatics Long-term Support project of the National Bioinformatics Infrastructure Sweden to Alexander Suh and Francisco J. Ruiz-Ruano. All the zebra finches sampled in this work were housed in aviaries through funds from the Max Planck Society to Bart Kempenaers. The computations were enabled by resources provided by the National Academic Infrastructure for Supercomputing in Sweden (NAISS) at UPPMAX, funded by the Swedish Research Council through grant agreement no. 2022-06725.

## Author contributions

Conceptualization: N.V., A.S.; Supervision: A.S., F.J.R.R., S.I.; Logistics: all authors; Analysis: N.V.; Visualization: N.V., Y.P.; Writing – original draft: N.V. with input from A.S.; Writing – review & editing of the second draft: N.V. with input from A.S., F.J.R.R., S.I.; Writing – review & editing of the final draft: N.V. with input from all authors.

